# Effects of bilateral and unilateral mental practice on actual motor performance and event-related desynchronization

**DOI:** 10.1101/2025.05.22.655524

**Authors:** Kazuya Umeno, Yoshihiro Itaguchi

## Abstract

Bilateral hand movement has been reported to enhance actual motor performance and increase motor cortex activation more than unilateral hand movement. This study investigated the interaction effects of bilateral and unilateral mental practice on event-related desynchronization (ERD) amplitude and unilateral motor performance, to explore better clinical rehabilitation for stroke patients with hemiparesis. The task was a ball rotation, where participants circulated two balls on their palm by using their fingers. Seventy-six healthy young adults were randomly assigned to one of three mental practice groups or a control group without mental practice. In the mental practice groups, three subgroups of participants engaged in ball-rotation motor imagery for two blocks in different orders: bilateral hand imagery followed by left hand imagery (BL group), left hand imagery followed by bilateral hand imagery (LB group), and consecutive left hand imagery (LL group). The results showed greater improvement in actual motor performance in the BL group than in the control group, while the performance of the LB and LL groups did not significantly differ from that of the control group. Greater ERD was observed in the BL group than in the LL group in the early sets of the first block. However, ERD amplitude decreased with continued mental practice, and significant correlations between ERD and motor performance were limited. These results suggest that engaging in bilateral mental practice before unilateral mental practice is beneficial in improving unilateral motor performance. Bilateral mental practice following unilateral mental practice may provide neither notable improvements nor detrimental effects on unilateral motor action. Our findings also reveal a mismatch between ERD and motor performance, implying that these two measures are not straightforwardly linked.

## Introduction

Motor imagery has been shown to be effective in enhancing actual motor performance. Previous studies have demonstrated its effectiveness in improving various motor tasks, including pinky finger abduction, elbow flexion, dart throwing, and toe abduction in healthy subjects [1-3]. Moreover, it enhances upper limb motor function and daily activity performance in stroke patients [4]. The National Clinical Guideline for Stroke recommends mental practice as a cost-effective and easily applicable intervention for upper limb rehabilitation in this population [5]. In this study, we define motor imagery as the mental simulation of movement, and mental practice as repeated motor imagery used as a training method to improve motor performance. Motor imagery activates brain areas involved in actual movement execution. For example, imagining simple finger movements activates cortical regions contralateral to the imagined hand, including the inferior parietal cortex, prefrontal cortex, anterior cingulate cortex, premotor cortex, and dorsolateral prefrontal cortex [6]. Hetu et al. [7] reported that motor imagery activates frontal-parietal, subcortical, and cerebellar regions involved in actual movement. Similarly, Guillot et al. [8] found overlapping activation in motor-related brain areas, including the inferior and superior parietal lobules, during both actual movement and motor imagery.

Bilateral hand movements have been shown to enhance actual motor performance, providing insight into the design of mental practice protocols. Previous studies have demonstrated that bilateral hand movements result in better performance compared to unilateral hand movements [9, 10]. Naito et al. [11] demonstrated that healthy older adults who performed bilateral hand movements exhibited greater improvements than unilateral hand movements in a pegboard task. Furthermore, the study found that improvements in the bilateral practice group were positively correlated with decreased excitability of the ipsilateral motor cortex (interhemispheric inhibition) observed using functional magnetic resonance imaging (fMRI) [11]. In stroke patients with chronic hemiplegia, Cunningham et al. [12] reported that bilateral elbow extension exercises led to greater improvements in the movement of the affected arm than unilateral exercises.

The effects of bilateral hand motor imagery on the improvement of actual motor performance remain poorly understood. Levin et al. [13] investigated changes in corticospinal excitability during motor imagery of bilateral wrist flexion and extension using transcranial magnetic stimulation. Their findings indicated that bilateral hand motor imagery induced larger motor evoked potentials (MEPs) compared to unilateral motor imagery. Additionally, an fMRI study reported that bilateral hand motor imagery enhanced functional connectivity between motor and sensory areas [14]. These findings suggest that bilateral hand motor imagery modulates movement-related brain activity, although its effects on improving actual motor performance remain inconclusive.

The combination and order of bilateral and unilateral hand movements may influence improvements in actual motor performance. Smith et al. [15] investigated wrist flexion and extension and found that bilateral hand movements enhanced subsequent performance of unilateral hand movements. Nozaki et al. [16] examined reaching tasks and reported that combining unilateral and bilateral hand movement learning produced greater learning effects than unilateral learning alone. Moreover, Hayashi [17] reported a transfer of learning from bilateral to unilateral hand movements, suggesting that combining these movement types may enhance performance. Consistent with these findings, several studies have shown that combining different types of training is more effective in enhancing both motor performance and brain activity than repeating practice of a single training type [18-20]. These findings indicate that interaction effects of bilateral and unilateral hand movements may enhance both actual motor performance and associated brain activity.

In summary, this study investigated how differences in the hand used for mental practice, as well as their combination, affect changes in actual motor performance and brain activity. Many studies on event-related desynchronization (ERD) feedback rely on classification accuracy derived from ERD amplitude and typically report only pre- and post-intervention data or representative cases [21-27]. A more detailed understanding of how ERD changes over the course of practice may enhance the learning effects of ERD feedback in both stroke patients and healthy subjects. Accordingly, we examined the temporal dynamics of ERD amplitude induced by neurofeedback training. We specifically hypothesized that the group engaging in both bilateral and unilateral mental practice would exhibit greater improvements in both actual motor performance and brain activity than the group practicing only unilateral imagery.

## Materials and methods

### Participants

Seventy-six healthy young adults (32 males and 44 females; mean age = 20.0 ± 1.4 years) participated in this study. They were randomly assigned to either a mental practice group or a control group. The mental practice group was further divided into three subgroups based on the order of hand use during imagery: the BL group (bilateral hand imagery followed by left hand imagery; 9 males and 13 females; mean age = 19.8 ± 1.3 years), the LB group (left hand imagery followed by bilateral hand imagery; 9 males and 13 females; mean age = 20.4 ± 1.4 years), and the LL group (consecutive left hand imagery; 11 males and 11 females; mean age = 20.0 ± 1.4 years). The control group (3 males and 7 females; mean age = 19.3 ± 1.0 years) did not engage in mental practice. The required sample size was calculated using G*Power 3 software. Based on a previous study [13], the effect size was set at 0.4, the significance level at 0.05, and the statistical power at 0.8. None of the participants had previous experience with the ball rotation task or had any serious orthopedic or central nervous system disorders affecting the upper extremities. All participants were right handed according to Chapman’s handedness questionnaire [28]. Written informed consent was obtained from all participants prior to the experiment. This study was approved by the Ethics Committee of the Tokoha University (No. 22-05). The study was conducted between January 2023 and March 2024, during which participants were recruited and data were collected.

### Mental Practice

In this study, mental practice was implemented using a ball rotation task based on the method described by Nojima [29] (Fig 1a). Participants sat in a relaxed position with both arms placed on a desk and circulated two balls on their palm as quickly as possible (Fig 1b). The ball was rotated clockwise with the right hand and counterclockwise with the left hand. Task instructions were presented on a monitor placed in front of the participants.

**Fig 1.**
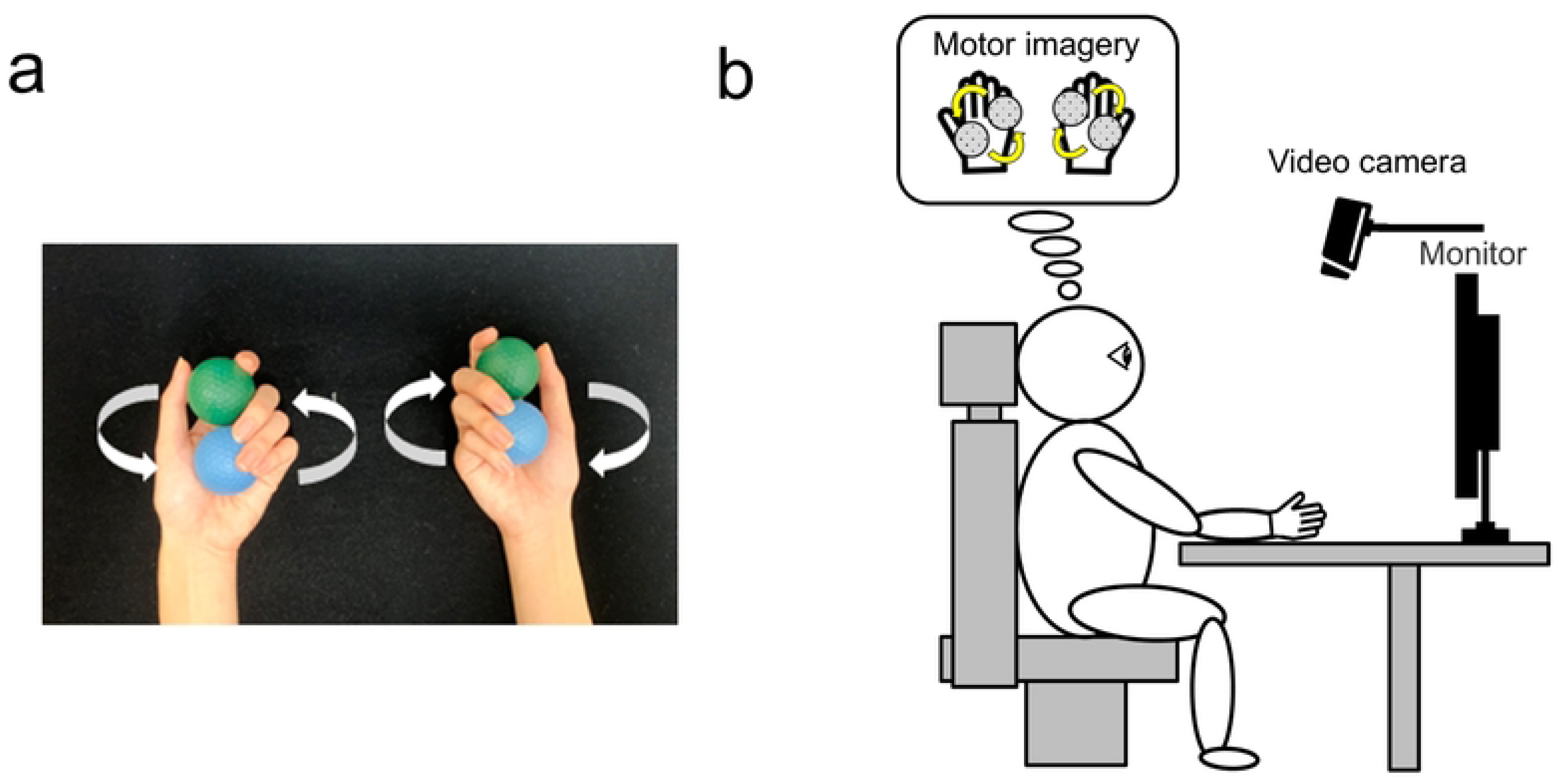
Experiment setting. (a) The ball rotation task for the bilateral hand. The ball was circulated clockwise with the right hand and counterclockwise with the left hand. (b) The experimental setup for the ball rotation task. Participants were seated on a chair with both arms placed on a desk. Video recordings were captured from above the participants’ hands. Task instructions were presented on a monitor positioned in front of the participant. In the motor imagery sessions, participants simply imagined that they were performing the task without moving their hands.

Two mental practice sessions were conducted, each lasting approximately 15 minutes, and motor performance was assessed before and after the sessions (pre test, test 1, and test 2) (Fig 2a). Each mental practice session consisted of two blocks, with each block comprising eight sets. In each set, participants performed 10 trials for each of the two imagery sessions: left hand imagery or bilateral hand imagery (10 trials × 8 sets) (Fig 2a). Each trial consisted of a baseline period, a motor imagery period (2 s), and visual feedback period (2 s) (Fig 2b). Video recordings were taken from above the participants’ hands during the task, and the number of rotations was counted offline.

**Fig 2.**
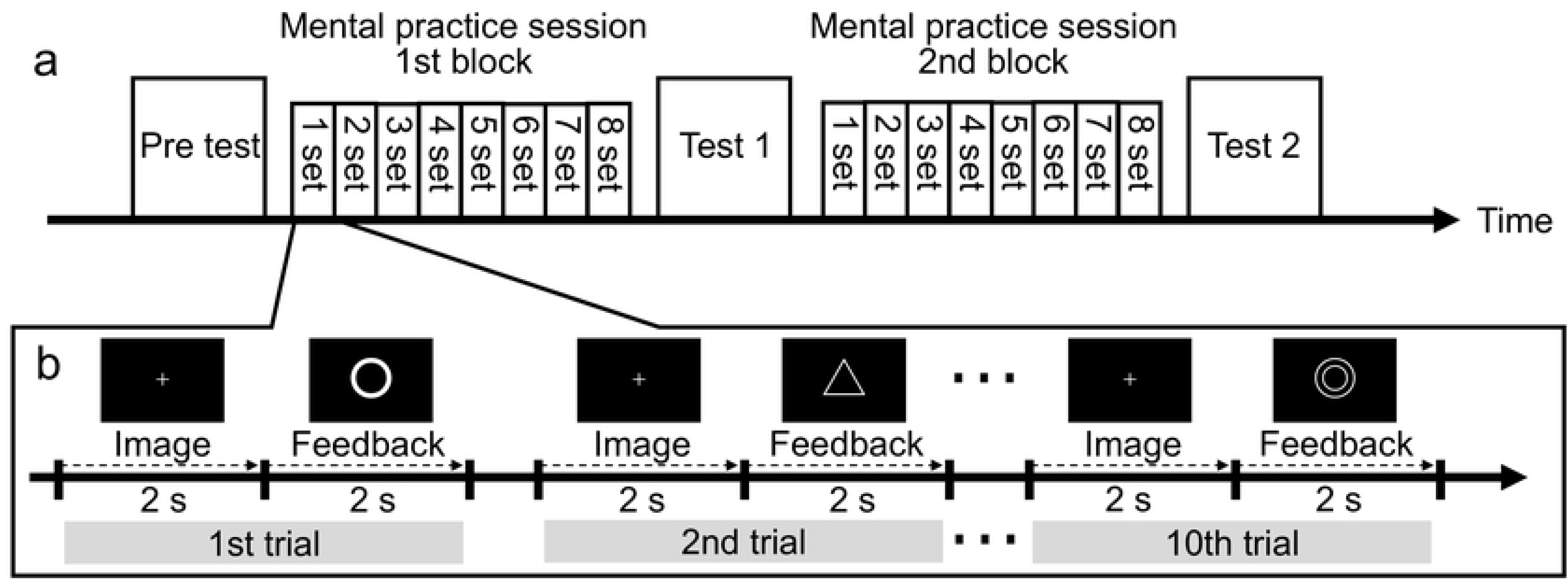
Experimental procedure. (a) Two mental practice sessions were conducted (each lasting approximately 15 minutes), and actual motor performance was assessed before and after the sessions (pre test, test 1, and test 2). (b) Time course of a trial and visual feedback provided to participants.

To evaluate motor performance improvement in the ball rotation task, we defined improvement scores based on the differences in the number of rotations between the test phases: improvement score after the 1st block represented the difference between the pre test and test 1, and improvement score after the 2nd block represented the difference between test 1 and test 2.

### EEG measurements

Brain activity during mental practice was assessed by recording electroencephalography (EEG) signals from four electrodes (F_P1_, F_P2_, C_3_, and C_4_) using gold cup electrodes connected to an OpenBCI Cyton Biosensing Board (OpenBCI). The reference and ground electrodes were placed on the left and right earlobes, respectively. The EEG signals were sampled at 250 Hz. The signals were bandpass filtered between 8 and 13 Hz. Video was recorded during the experiment to ensure that participants kept their hands still during the mental practice. EEG analyses were conducted using MATLAB R2020a (MathWorks, Inc.). For EEG signal processing, data exceeding three standard deviations from the baseline amplitude were excluded as noise. EEG segments with amplitudes exceeding three standard deviations from the baseline period were removed to eliminate noise.

ERD was calculated using a method proposed by Pfurtscheller [30]. Alpha ERD recorded over motor related regions is commonly used in motor imagery tasks and brain– machine interface systems [31, 32]. ERD was defined as the percentage decrease in alpha amplitude during motor imagery relative to the baseline value. It was calculated by dividing the difference between the root mean square amplitude of the alpha wave during the imagery period (A) and that during the baseline period (R) by the root mean square amplitude during the baseline period (R) (Equation 1).

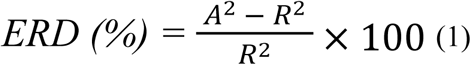

### Neurofeedback

A neurofeedback system based on EEG signals was developed and used in this study. EEG signals were transmitted from the OpenBCI device to MATLAB using the BrainFlow MATLAB package. ERD was calculated in real time in MATLAB and visually displayed on a monitor as feedback. ERD was calculated from the C_4_ electrode (above the right motor cortex) for left hand motor imagery, and from the average of C_3_ and C_4_ electrodes for bilateral hand motor imagery. Participants were seated in a chair with both arms placed on a desk and were instructed to fixate on a monitor. During both the motor imagery and baseline periods, a fixation point was displayed, and participants were instructed to avoid blinking. After each imagery period (2 s), visual feedback was provided based on the magnitude of the calculated ERD. In this study, greater ERD was represented by more negative values. The following visual symbols were used to represent ERD: a double circle (◎) for ERD less than −20%, a circle (〇) for ERD between −10% and −20%, a triangle (△) for ERD between 0% and −10%, and a minus sign (−) for ERD greater than 0% (Fig 2).

### Statistical analysis

Three statistical analyses were conducted. First, to examine the effects of mental practice using bilateral hand movements on actual motor performance, a three-way repeated measures ANOVA (group × hand × block) was performed on the changes in the number of ball rotations. Second, to examine the effects of the hand used during mental practice on ERD amplitude, a two-way repeated measures ANOVA (group × set) was performed on the ERD. Last, to examine the relationship between the magnitude of ERD and improvements in actual motor performance, Spearman’s rank correlation was used to assess the association between changes in the number of ball rotations and the average ERD for each set. All multiple comparisons were corrected using Holm’s method. Statistical analyses were performed using R software (version 4.2.0).

## Results

### Actual motor performance in 1st-and 2nd-blocks

Fig 3 shows the changes in the number of ball rotations during the 1st and 2nd blocks relative to the baseline period. For unilateral (left hand) motor performance, the mean changes were 4.8 (*SD* = 2.2) in the BL group, 3.8 (*SD* = 1.9) in the LB group, 4.4 (*SD* = 2.1) in the LL group, and 2.5 (*SD* = 1.5) in the control group. A three-way repeated measures ANOVA (group × hand × block) on the changes in the number of ball rotations revealed significant main effects of group (*F*(3, 72) = 3.47, *p* = 0.020, η_p_² = 0.13) and block (*F*(1, 3) = 9.46, *p* = 0.003, η_p_² = 0.12). For bilateral motor performance, the mean changes in the number of rotations across the two blocks were 5.1 (*SD* = 2.0) in the BL group, 3.6 (*SD* = 1.9) in the LB group, 4.3 (*SD* = 2.1) in the LL group, and 3.1 (*SD* = 1.3) in the control group. Additionally, multiple comparisons revealed significant differences between the BL group and the control group in the number of ball rotations (*t*(72) = 2.9, *p* < 0.05).

**Fig 3.**
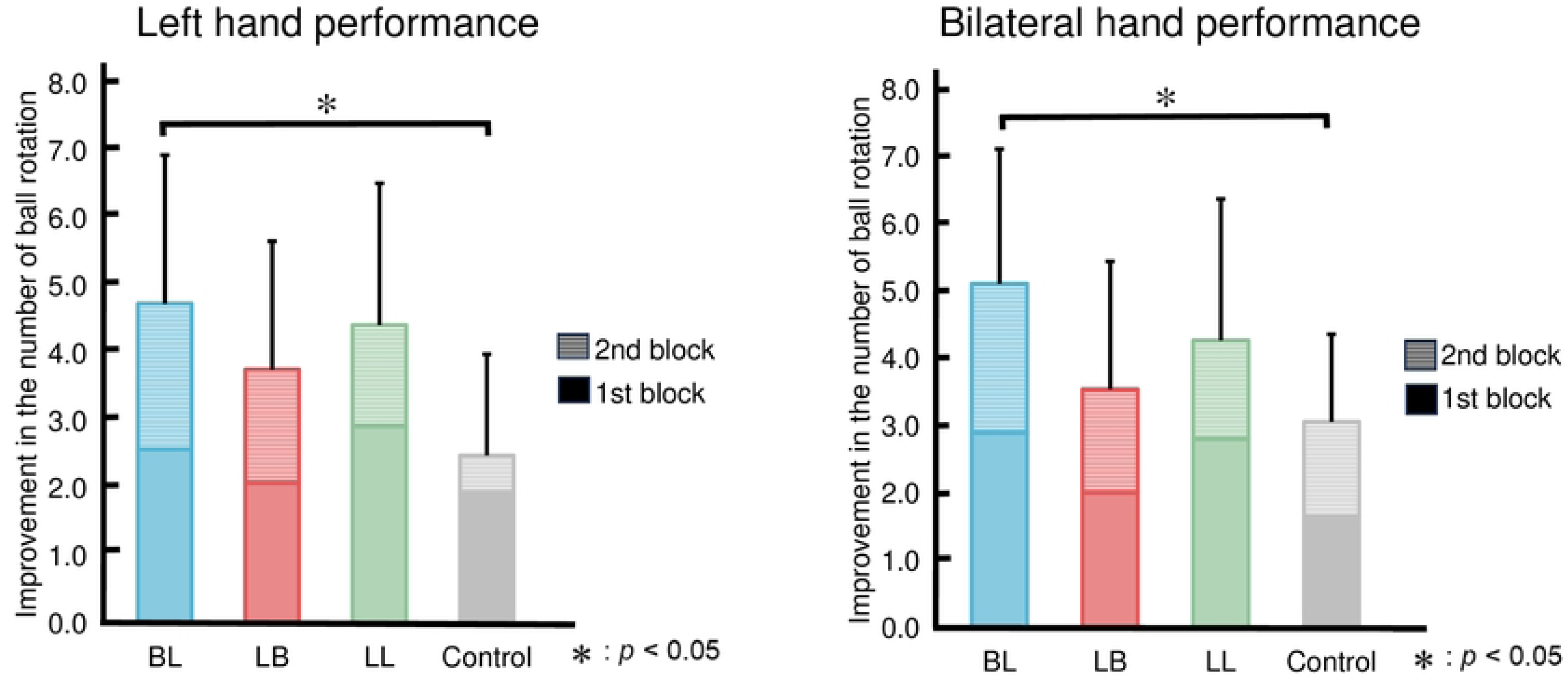
Changes in motor performance for each group (1st block + 2nd block) The vertical axis represents the change in the number of ball rotations. Performance improvement was significantly greater in the BL group compared to the control group.

### ERD analysis

Mean ERD values were calculated by averaging the ERD from 10 trials into one set, and then averaging these set values across all participants in each group. In the 1st block, the mean ERD values were −15.9% (*SD* = 11.6) at C_3_ and −15.7% (*SD* = 15.0) at C_4_ in the BL group, −11.2% (*SD* = 9.9) at C_3_ and −13.7% (*SD* = 12.0) at C_4_ in the LB group, and −10.6% (*SD* = 14.2) at C_3_ and −10.0% (*SD* = 13.0) at C_4_ in the LL group. In the 2nd block, the mean ERD values were −13.1% (*SD* = 12.3) at C_3_ and −14.0% (*SD* = 17.3) at C_4_ in the BL group, −10.2% (*SD* = 10.1) at C_3_ and −12.1% (*SD* = 11.9) at C_4_ in the LB group, and −9.8% (*SD* = 14.6) at C_3_ and −8.0% (*SD* = 13.5) at C_4_ in the LL group (Table 1).

**Table 1.** Mean event-related desynchronization per block for each group.

| Group | Electrode | Event-Related Desynchronization (%) |  |
| --- | --- | --- | --- |
|  |  | 1st block | 2nd block |
| BL | $C_3$ | -15.9 (11.6) | -13.1 (12.3) |
| | $C_4$ | -15.7 (15.0) | -14.0 (17.3) |
| LB | C<sub>3</sub> | -11.2 (9.9) | -10.2 (10.1) |
|  | C<sub>4</sub> | -13.7 (12.0) | -12.1 (11.9) |
| LL | C<sub>3</sub> | -10.6 (14.2) | -9.8 (14.6) |
|  | C<sub>4</sub> | -10.0 (13.0) | -8.0 (13.5) |

Values represent the mean ERD (%) with standard deviations in parentheses. Negative values indicate the presence of event-related desynchronization. BL: Bilateral hand motor imagery in the 1st block and left hand motor imagery in the 2nd block. LB: Left hand motor imagery in the 1st block and bilateral hand motor imagery in the 2nd block. LL: Left hand motor imagery in both the 1st and 2nd blocks

The time series data of ERD across the 8 sets within each block were visually examined prior to conducting statistical analyses on the quantitative data. Fig 4 shows the time course of ERD across the 8 sets within each block for the three groups that engaged in mental practice. In this figure, both the BL and LB groups exhibited the largest ERD in the 1st or 2nd set of the 1st block. ERD gradually decreased as mental practice progressed, and this change was more pronounced in the BL group. Additionally, ERD increased in the 1st set of the 2nd block following actual motor performance.

**Fig 4.**
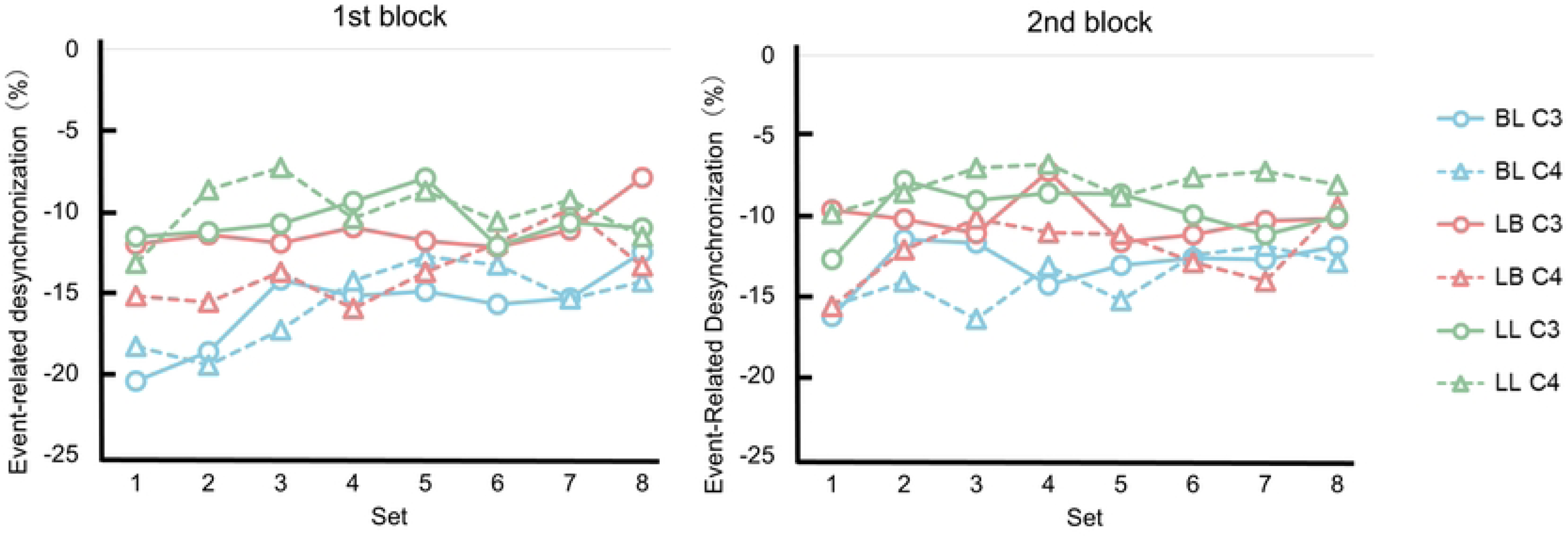
Changes in event-related desynchronization across sets for each group. The vertical axis represents event-related desynchronization, and the horizontal axis represents the set number within each mental practice session.

A two-way ANOVA (group × set) revealed a significant main effect of set on C_3_ in the 1st block (*F*(14, 441) = 2.59, *p* = 0.012, η_p_² = 0.04). No significant main effects were observed in the 2nd block. Multiple comparisons revealed no significant differences between groups in either the 1st or 2nd block. For C_4,_ a main effect of set was observed in the 1st block (*F*(7, 441) = 2.77, *p* = 0.008, η_p_² = 0.04). Although the interaction between group and set was not statistically significant, a trend toward significance was observed (*F*(14, 441) = 1.70, *p* = 0.053, η_p_² = 0.05). A simple main effects analysis revealed significant group differences at the 2nd and 3rd sets (2nd set: *F*(2, 63) = 3.20, *p* = 0.047; 3rd set: *F*(2, 63) = 3.58, *p* = 0.034). ERD in the BL group was lower than in the LL group in the 2nd and 3rd sets, with significant differences observed (*p* = 0.045 and *p* = 0.031). Multiple comparisons revealed a significant difference between the 1st and 7th sets (*t*(63) = 3.37, *p* = 0.0013, adjusted *p* = 0.036). In the 2nd block, a significant main effect of set was observed (*F*(7, 441) = 2.06, *p* = 0.047, η_p_² = 0.03). Multiple comparisons revealed no significant differences between groups.

### Relationship between ERD amplitude and ball rotation performance

Fig 5 shows the correlations between ERD at the C_3_ and C_4_ electrodes and improvement scores in ball rotation performance in the BL, LB, and LL groups. In the BL group, negative correlations were observed between ERD and bilateral improvement score after the 1st block in ball rotation performance at both C_3_ and C_4_. Statistically significant correlations were found in three specific sets: at the C_3_ electrode during the 1st block, 8th set (*r* = −0.437, *p* = 0.041); at the C_4_ electrode during the 2nd block, 7th set (*r* = −0.466, *p* = 0.028); and 8th set (*r* = −0.476, *p* = 0.024) of the same block. In the LB group, weak positive correlations were observed between ERD and bilateral improvement score in the 1st block across both blocks. A statistically significant correlation was identified at the C_4_ electrode during the 1st block, 1st set (*r* = 0.443, *p* = 0.029). In the LL group, weak negative correlations were observed between ERD and improvement score in the 2nd block. Statistically significant correlations based on left improvement scores were found at the C_3_ during the 1st block in three consecutive sets: 2nd set (*r* = −0.431, *p* = 0.044), 3rd set (*r* = −0.435, *p* = 0.043), and 4th set (*r* = −0.433, *p* = 0.043). In addition, a significant correlation based on bilateral improvement scores was found at the C_4_ electrode during the 2nd block, 7th set (*r* = −0.462, *p* = 0.030). Overall, across the three mental practice groups, the correlations between ERD at C_3_ and C_4_ and improvement scores in ball rotation performance were generally weak.

**Fig 5.**
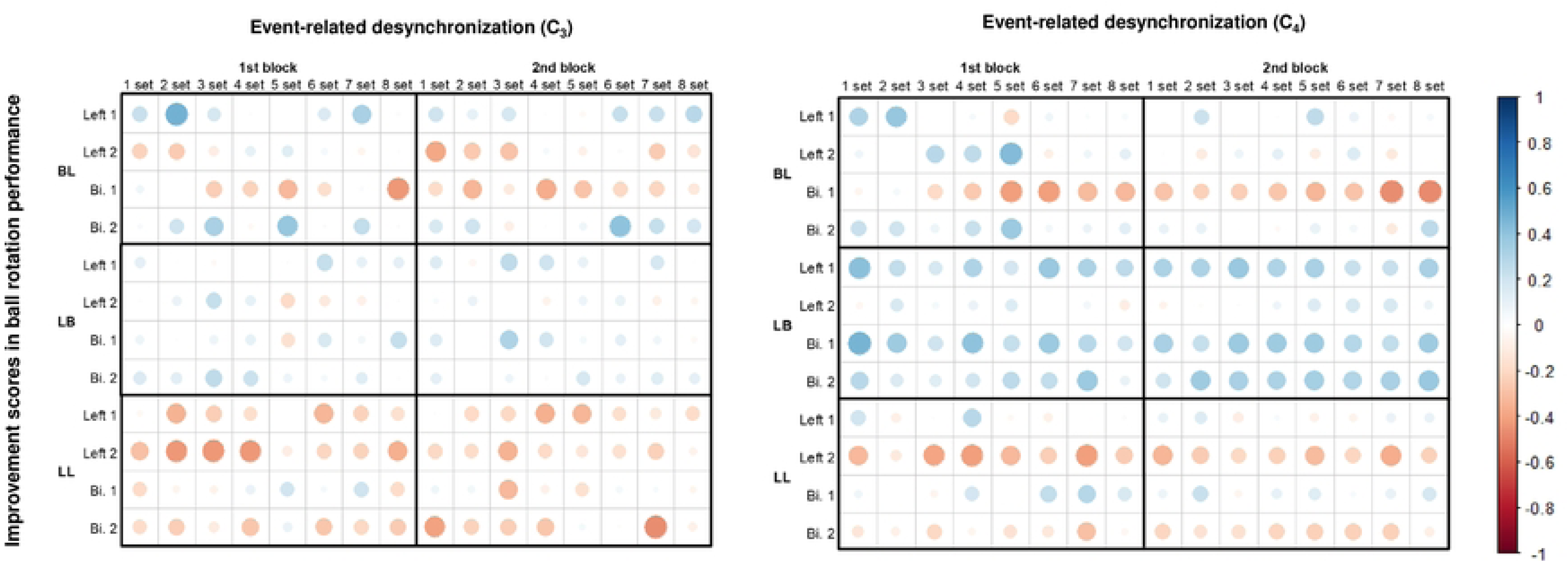
Correlation coefficients between event-related desynchronization amplitudes and improvement scores in ball rotation performance. Red circles indicate negative correlations and blue circles indicate positive correlations. The intensity and size of the circle represent the strength of the correlation. Most of the correlations were not statistically significant. Bi: bilateral hands.

## Discussion

This study investigated the effects of different hand use and their combinations in mental practice on actual motor performance and brain activity. We hypothesized that engaging in bilateral hand motor imagery followed by left hand imagery would lead to greater improvement in ball rotation performance than performing left hand imagery alone. The results showed that only the BL group exhibited a significantly greater increase in the number of ball rotations compared to the control group, supporting our hypothesis. These results suggest that engaging in bilateral hand imagery prior to left hand imagery may be effective in improving actual motor performance. We also hypothesized that engaging in both bilateral and unilateral imagery would result in greater ERD compared to unilateral imagery alone. The results showed a significant difference between the BL and LL groups in the early sets of the 1st block, which does not fully support our hypothesis. This finding suggests that bilateral hand imagery, at least during the early phase of the task, may be more effective in enhancing brain activity than left hand imagery. To our knowledge, this is the first study to report that improvements in actual motor performance and changes in ERD may not fully correspond when assessed using both behavioral and neural measures.

### Improvement in actual motor performance

Engaging in bilateral mental practice followed by unilateral mental practice may be effective in improving actual motor performance. In the present study, the BL group demonstrated significantly greater improvement in motor performance compared to the control group. Nozaki et al. [16] reported that partial transfer of learning occurs between unilateral and bilateral hand movements during actual motor tasks. Torres et al. [18] showed that combining different types of practice led to greater improvement in motor performance than repeating the same type of practice. Based on these findings, the transfer effect of engaging in both bilateral and unilateral mental practice may have contributed to the observed improvement in motor performance in the present study. Thus, our results suggest that such a combination may facilitate motor performance, similar to actual motor execution.

The effect of the order of mental practice on motor performance improvement has not been thoroughly examined in previous studies. In the present study, motor performance improved in the BL group, whereas the improvement in the LB group was limited. This difference suggests that the order of imagery in mental practice may influence performance. Smith et al. [15] reported that motor performance improved when bilateral hand movements were followed by unilateral hand movements. These results indicate that the order of movements in mental practice may affect motor performance, similar to actual motor execution. Previous studies have reported that movement order is important in post stroke rehabilitation, with greater improvement observed when the unaffected hand is trained before the paretic hand [33, 34]. As mental practice has been widely used in stroke rehabilitation [4, 35, 36], the sequence of imagery tasks may need to be carefully considered in clinical applications.

### The effect of bilateral and unilateral mental practice on ERD

Bilateral mental practice may induce greater brain activity than imagery involving only the left hand. In the present study, the BL group exhibited greater ERD at C_4_ than the LL group during the 2nd and 3rd sets of the 1st block. This result supports our previous findings [37], which demonstrated that bilateral mental practice induced greater ERD than left hand imagery, although inconsistencies emerged in the later phase. These inconsistencies may stem from differences in task design; the current study employed a block design, whereas the previous study alternated the hand used during mental practice. Levin et al. [13] compared MEPs between unilateral and bilateral hand motor imagery and reported that bilateral hand imagery induced greater corticospinal excitability. Although the previous study used MEPs and the present study used ERD to evaluate brain activity, both measures consistently showed that bilateral hand motor imagery leads to stronger brain activation. ERD is considered to reflect brain activity related to motor execution, motor imagery, or action observation [38], and a greater ERD amplitude reflects stronger activation of the primary motor cortex [39]. Previous research has also shown that bilateral hand motor imagery enhances corticospinal excitability more than unilateral imagery, supporting our finding that ERD was greater during bilateral hand imagery. Thus, the greater ERD observed in the bilateral hand imagery in this study is consistent with existing literature.

Continued mental practice may lead to a decrease in ERD amplitude. Vasilyev et al. [40] conducted continued mental practice combined with EEG-based neurofeedback and reported that stable ERD was observed. In contrast, the present study demonstrated a decrease in ERD amplitude with continued mental practice, which is inconsistent with previous findings. We propose two possible explanations for this result. First, continued mental practice of a single movement may have induced fatigue. Previous studies have shown that fatigue increases alpha power [41-43], and this increase may account for the reduction in ERD observed in the present study. Second, the smaller ERD may reflect a decrease in baseline alpha power. Several studies have reported that performing cognitive task reduces resting alpha power [44-46]. ERD is defined as a reduction in alpha power associated with a specific event, typically calculated as the percentage decrease from baseline alpha amplitude [30]. In the present study, continued mental practice may have reduced the baseline alpha amplitude, which served as the reference for ERD calculation, thereby resulting in smaller ERD amplitude. The mechanisms underlying ERD reduction have not been fully explored, and its cause remains unclear, warranting further investigation.

### Relationship between brain activity and actual motor performance

Mental practice may enhance the magnitude of ERD, which may not necessarily align with improvement in actual motor performance. Previous studies using PET and fMRI have reported that greater brain activity during physical practice is associated with better actual motor performance [47, 48]. Based on these findings, the present study hypothesized that the magnitude of ERD during mental practice would be significantly correlated with changes in actual motor performance. Although actual motor performance improved following mental practice, ERD amplitude did not continue to increase, and no significant correlation was found between ERD amplitude and actual motor performance improvement. This discrepancy may be attributed to the differing measurement characteristics of EEG compared to neuroimaging techniques such as PET and fMRI.

PET and fMRI assess brain activity by detecting metabolic changes related to cerebral blood flow, making them suitable for observing gradual changes in brain activation during continuous tasks [49, 50]. In contrast, EEG offers high temporal resolution and captures transient changes in neural activity [51], which may reduce its sensitivity to gradual changes in brain activity compared to PET and fMRI [49]. These methodological differences may explain the inconsistency between ERD changes and actual motor performance improvements. Although no previous studies have directly demonstrated a relationship between increased ERD amplitude and improved actual motor performance, both our previous study [37] and the present study consistently demonstrated this dissociation.

### Limitation of the Study

The present study has several limitations. First, similar to many previous studies on mental practice [52-55], it was limited to healthy adults and employed a specific motor imagery task. Future studies should investigate how mental practice affects actual motor performance and brain activity by including different participant characteristics and varying types of motor imagery. It may be beneficial to conduct a similar study involving older adults and stroke patients to determine whether comparable patterns of brain activity and actual motor performance can be observed. The current findings may provide a basis for further investigation into the relationship between mental practice and brain activity, and future studies may include more in-depth EEG analyses. Additionally, this study focused on specific outcomes of mental practice, namely ERD and improvement in actual motor performance. Complementary assessments of other factors related to mental practice, such as EEG frequency bands other than alpha [56], fatigue [57], and the quality of motor imagery [58] may help refine the current understanding and clarify the mechanisms underlying the interaction between brain activity and actual motor performance.

## Conclusion

The present study demonstrated the effects of bilateral mental practice on actual motor performance. In particular, the combination of bilateral and unilateral mental practice contributed to improvements in actual motor performance, suggesting that this approach may hold potential as a rehabilitation strategy. Additionally, ERD amplitude decreased with continued mental practice. Although the underlying cause of this decrease remains unclear, it may be related to fatigue or a reduction in baseline alpha amplitude. The findings of this study suggest a dissociation between improvements in actual motor performance and changes in ERD amplitude with continued mental practice, which has not been clearly reported in previous studies. Further research is needed to elucidate how changes in ERD amplitude affect actual motor performance and to explore the clinical applicability of mental practice protocols informed by neural activity.

## Disclosure statement

No potential conflict of interest was reported by the author (s).

## Funding

This study was supported by the Japan Society for the Promotion of Science, KAKENHI (grant number 21K11301).

## Acknowledgments

The authors are grateful to the students at Tokoha University for their help with the data acquisition.

## Author Contributions

Conceptualization: Kazuya Umeno, Yoshihiro Itaguchi.

Data curation: Kazuya Umeno.

Formal analysis: Kazuya Umeno.

Funding acquisition: Kazuya Umeno.

Investigation: Kazuya Umeno, Yoshihiro Itaguchi.

Methodology: Kazuya Umeno, Yoshihiro Itaguchi.

Project administration: Yoshihiro Itaguchi.

Resources: Kazuya Umeno, Yoshihiro Itaguchi.

Software: Kazuya Umeno.

Supervision: Yoshihiro Itaguchi.

Validation: Kazuya Umeno, Yoshihiro Itaguchi.

Visualization: Kazuya Umeno, Yoshihiro Itaguchi.

Writing – original draft: Kazuya Umeno.

Writing – review & editing: Kazuya Umeno, Yoshihiro Itaguchi.

## Notes

### Competing Interest Statement

The authors have declared no competing interest.

